# *Helicobacter pylori* infection and α-synuclein pathology drive parallel neurodegenerative pathways in the substantia nigra

**DOI:** 10.1101/2025.08.26.672190

**Authors:** Alejandro Soto-Avellaneda, Alice Prigent, Lindsay Meyerdirk, Noah Schautz, John Andrew Pospisilik, Lena Brundin, Michael X. Henderson

**Affiliations:** Department of Neurodegenerative Science, Van Andel Institute, Grand Rapids, MI 49503; Department of Epigenetics, Van Andel Institute, Grand Rapids, MI 49503

**Keywords:** Inflammation, T cells, infection, Parkinson’s disease, *SNCA*, *Helicobacter pylori*

## Abstract

Parkinson’s disease is a common neurodegenerative disease related to both genetic and environmental insults. Epidemiological studies have linked *Helicobacter pylori (H. pylori*) infection to Parkinson’s disease risk, but the underlying mechanisms of this association remain unclear. In this study, we investigate whether chronic infection with a pathogenic *H. pylori* strain can induce α-synuclein aggregation or neurodegeneration, and whether infection clearance mitigates these effects. We also assessed whether *H. pylori* infection exacerbates α-synuclein pathology and neuron loss when combined with seeding of α-synuclein pathology. We find that chronic *H. pylori* infection induces a sustained immune response in the gut and plasma that leads to mild brain inflammation and dopaminergic neuron loss, independent of α-synuclein pathology. These effects are attenuated by eradication of the infection. In mice with α-synuclein pathology induced by pre-formed fibrils, *H. pylori* did not further exacerbate the extent of pathology or neuronal death. Together, these results suggest that *H. pylori* infection can lead to neurodegeneration through inflammatory mechanisms independent of α-synuclein aggregation. Our findings offer mechanistic insights into how pathogens influence the risk and progression of Parkinson’s disease.

## NTRODUCTION

Parkinson’s disease (PD) is a progressive neurodegenerative disorder characterized by the loss of dopaminergic neurons in the substantia nigra pars compacta and the accumulation of Lewy pathology composed largely of misfolded α-synuclein^1,2^. While PD is classically defined by its motor symptoms, it has a prolonged prodromal phase in which non-motor symptoms, including constipation and gut dysbiosis, frequently appear years before diagnosis^3,4^. The presence of early α-synuclein pathology in the olfactory bulb and dorsal motor nucleus of the vagus has led to the hypothesis that environmental triggers in the gut may initiate disease^5-10^. In line with this hypothesis, enteric neurons and peripheral nerves display α-synuclein pathology in up to 52% of patients with PD^8,11,12^ and α-synuclein identified in skin peripheral nerves can be used to confirm a diagnosis for central α-synucleinopathies^13-16^.

Although approximately 10% of PD cases are attributable to known mutations in genes such as *SNCA, LRRK2, PINK1, and PRKN*^17^, the majority are idiopathic, with unclear initiation factors. Among the proposed environmental contributors to PD risk, infectious agents appear to play an important, although likely modulatory, role. Epidemiological studies have associated various viral and bacterial infections, including influenza, hepatitis C, Epstein-Barr, and *Helicobacter pylori (H. pylori)* with increased risk and severity of PD^18^. Several gut bacteria have also been associated with development of PD patients, including overgrowth of *Verrucomicrobiales, Lactobacillus, Bifidobacterium*, and *Helicobacter pylori* (*H. pylori*) infection^19,20^. These associations suggest that infection-induced inflammation or immune responses may play a role in PD onset or progression.

*H. pylori* is a stomach pathogen best known for its role in peptic ulcers and gastric cancer^21^, and its relevance to PD followed the initial observation of an increased occurrence of gastric ulcers in people with PD^22^. Since this initial association, seropositivity for *H. pylori* has been reported in 50% of people with PD and is associated with a poor response to levodopa, while eradication therapy has been linked to better clinical outcomes^23^. A bidirectional Mendelian randomization analysis suggested a causal relationship between *H. pylori* infection and PD motor and cognitive decline^24^. A second meta-analysis from 2020 found that among all infectious agents (bacterial, viral, fungal), *H. pylori* demonstrated the highest association to PD with an odds ratio of 1.6^25,26^. Despite strong epidemiological support, the biological mechanisms by which *H. pylori* may influence PD pathogenesis remain poorly understood, and to date it is unclear how *H. pylori* infection acts alone, or in concert with α-synuclein pathology, to impact PD-relevant pathways.

In this study, we address this gap by examining the impact of *H. pylori* infection in a mouse model. Specifically, we explored whether chronic infection with a known pathogenic strain of *H. pylori* could trigger systemic and neuroinflammatory responses, promote α-synuclein aggregation, or induce dopaminergic neuron loss. We find that *H. pylori triggers* a sustained immune response in the gut and plasma and leads to mild inflammation and neurodegeneration in the substantia nigra in the absence of α-synuclein pathology. These effects are mitigated by triple therapy eradication of *H. pylori*, though inflammation in the substantia nigra persists following *H. pylori* clearance. To evaluate whether *H. pylori* synergizes with α-synuclein pathology to worsen degeneration, we injected mice intracranially with α-synuclein pre-formed fibrils (PFFs) to induce seeded pathology. While α-synuclein PFF injection alone drives widespread α-synuclein pathology and neuron loss, *H. pylori* exposure did not further exacerbate these effects. Together, our results demonstrate that *H. pylori* infection can promote neurodegeneration through inflammatory processes independent of induction of α-synuclein pathology. These findings provide mechanistic insight into how pathogens may influence PD risk and progression.

## RESULTS

### *H. pylori* treatment leads to sustained infection that is reversible with triple therapy

To determine the effect of sustained *H. pylori* infection on PD-relevant phenotypes, we infected a group of mice with the pre-mouse Sydney strain 1 (PMSS1) of *H. pylori* ^27,28^ or vehicle at 3 months of age and allowed them to age to 9 months post-infection (**Fig. 1A**). This strain is known to chronically colonize the mouse pylorus. To differentiate between the effect of treated and sustained infections, we treated a second cohort of mice with *H. pylori* or vehicle followed by triple therapy consisting of omeprazole, amoxicillin, and clarithromycin^29,30^ one month after infection (**Fig. 1A**). *H. pylori* showed sustained colonization of the stomach across both 9-month infection cohorts as confirmed by immunofluorescence of gastric glands (**Fig. 1B**) and by PCR (**Fig. 1C, 1D**), while triple therapy was able to eradicate detectable *H. pylori* from the stomach (**Fig. 1D**).

**Figure 1.**
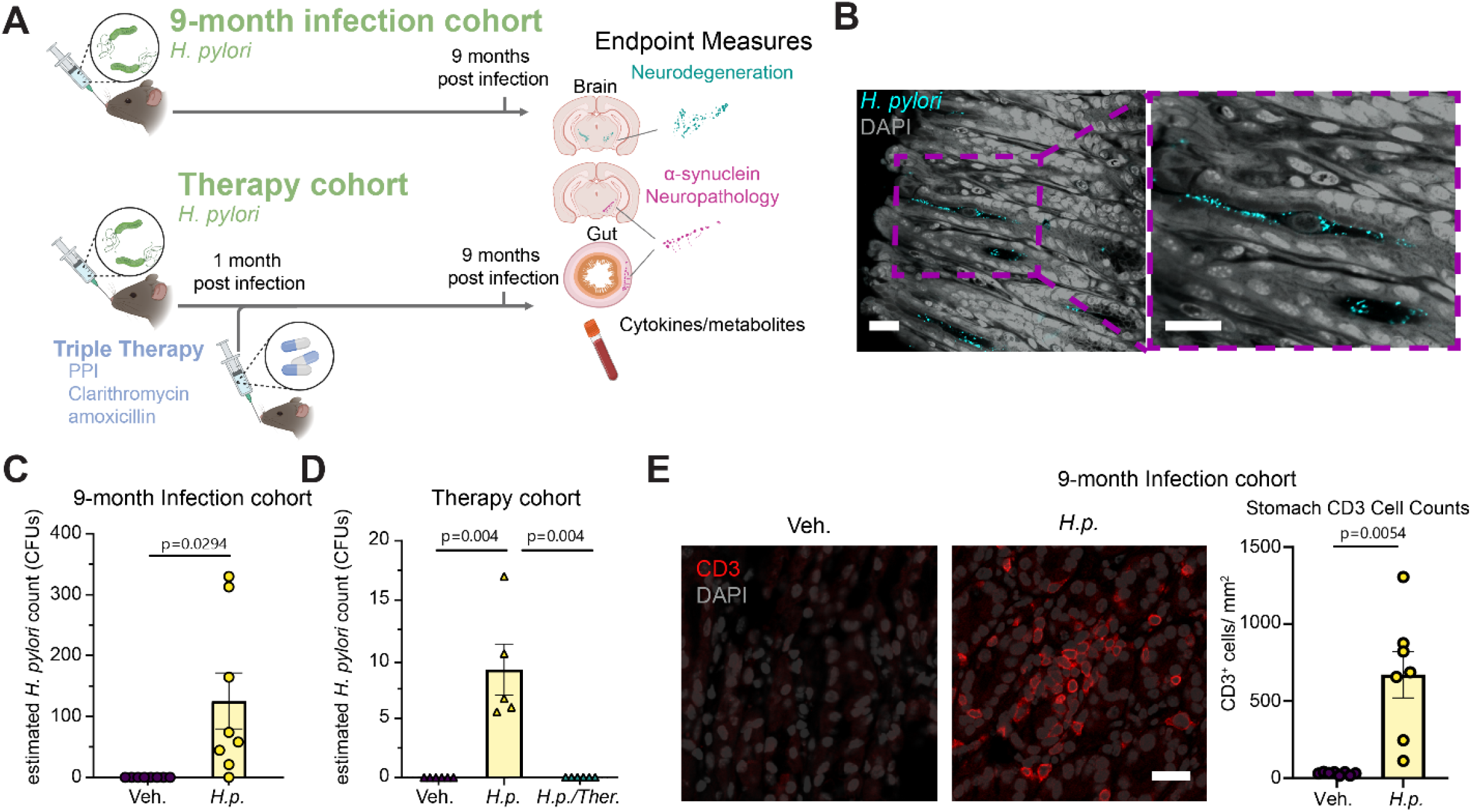
*H. pylori* treatment leads to sustained infection that is reversible with triple therapy. **A**. Experimental schematic. Mice were infected with *H. pylori* or fed DMEM vehicles. Mice were aged 9 months post initial infection. n = 8 (*H*.*p*), or 8 (Veh.) A second cohort of mice was infected with *H. pylori*. and a subset of the infected mice was treated with antibiotic triple therapy to clear the infection. n = 8 (Veh.), 6 (*H*.*p*.), or 8 (*H*.*p*.*/*Ther.) All mice were then aged for 9 months post-initial infection. Vehicle mice in the Therapy cohort also received triple antibiotic therapy. **B**. Representative image of *H. pylori*-infected stomach tissue stained for *H. pylori*. Scale bar = 20 µm. **C**. qPCR quantification of *H. pylori* load in 9-month Infection cohort and **D**. Therapy cohort *H. pylori* load. **E**. Representative image of 9-month Infection cohort mouse stomachs stained for CD3 and quantification of CD3-positive cells. A Welch’s T-test was performed for the 9-month infection cohort *H. pylori* load analysis, and to compare stomach CD3^+^ cell count. Scale bar= 20 µm. A one-way ANOVA with Tukey’s multiple comparisons test was performed for the Therapy cohort analysis. Statistical significance (p < 0.05) is indicated above relevant comparisons; all other differences are non-significant.

We hypothesized that *H. pylori* could lead to increased PD risk by inducing the aggregation of α-synuclein through an unknown mechanism. To test this hypothesis, we stained for phospho-serine 129 (pS129) α-synuclein throughout the brain of mice that had sustained 9 months of *H. pylori* infection. We were unable to detect α-synuclein aggregates in the brains of *H*.*pylori-*infected mice (**Fig. S1A**) while aggregates were detected in similar areas of positive control mice injected with α-synuclein PFFs (**Fig. S1B**). The dorsal motor nucleus of the vagus nerve (DMX) is an early site of α-synuclein aggregation in PD^31^ and is directly connected to the stomach through projections to the myenteric plexus. Therefore, were undertook a selective examination of cholinergic neurons within the DMX. Sections stained for choline acetyltransferase and pS129 α-synuclein revealed no α-synuclein inclusions in either vehicle or *H. pylori*-treated mice (**Fig. S1C**).

These data suggest that the increased risk conferred by *H. pylori* is not through direct induction of α-synuclein aggregation.

A second mechanism by which *H. pylori* could increase risk of PD is through inflammatory responses to infection that induce systemic inflammation or auto-immunity through molecular mimicry^32^. Yet, one of the reasons *H. pylori* is able to develop stable colonies is due to their release of immuno-suppressive peptides that prevent their clearance by the immune system^33^. To determine if 9 months of *H. pylori* colonization induced an immune response, we stained stomachs for CD3+ T-cells (**Fig. 1E**). *H. pylori* infection induced a substantial infiltration of CD3+ T-cells into the stomach compared to vehicle-treated controls (**Fig. 1E**).

### *H. pylori* infection induces local inflammation and systemic elevation of IFN-γ

Several cytokines are increased in PD cerebrospinal fluid and plasma, including tumor necrosis factor-α (TNF-α), interferon-γ (IFN-γ), interleukin-1β (IL-1β), and interleukin-6 (IL-6)^34-36^. Among these, TNF-α, IL-1β, and IL-6 are also increased in individuals with *H. pylori* infections^37^. Because of the overlap between the cytokines found in PD and *H. pylori* infection, we designed assays to determine if *H. pylori* induced local or systemic elevation of pro-inflammatory cytokines observed in PD.

We performed a multiplexed assay to assess the aforementioned cytokines as well as addition cyto- and chemokines IL-2, 4,5, 10, 12p70, and keratinocyte chemoattractant (KC/GRO) in stomach, blood plasma, and spinal cords of mice.

Mice treated with *H. pylori* showed elevation of IFN-γ, TNF-α, IL-1β, IL-2, and KC/GRO in the stomach, and triple therapy reversed this effect (**Fig. 2A-2D, S2A-S2F**). In plasma, IFN-γ and KC/GRO were the only molecules significantly elevated in the plasma of *H. pylori*-infected animals compared to vehicle control, although there was some discrepancy across cohorts (**Fig. 2E-2H, S2G-S2L**). Notably, in the plasma, several cytokines were near or below the detection limit. In spinal cord, none of the tested molecules were significantly elevated (**Fig. 2I-2L, S2M-S2O**). To further assess inflammation in the central nervous system, we conducted qPCR on spinal cords to determine whether there were RNA-level changes in *Ifng (*IFN-γ), *Il1b* (IL-1β), *Il6* (IL-6), *Il17a* (IL-17a), and *Tnf (*TNF-α). There was a slight elevation in average IFN-γ RNA (*Ifng*) in *H. pylori*-infected animal spinal cords, but no significant changes were observed in any cytokine RNAs (**Fig. S3**). Together, these experiments show that *H. pylori* infection leads to a sustained T-cell infiltration of the stomach and pro-inflammatory cytokine release. IFN-γ, reaches the plasma. IFN-γ was previously shown to be elevated to a similar extent in PD plasma^36^.

**Figure 2.**
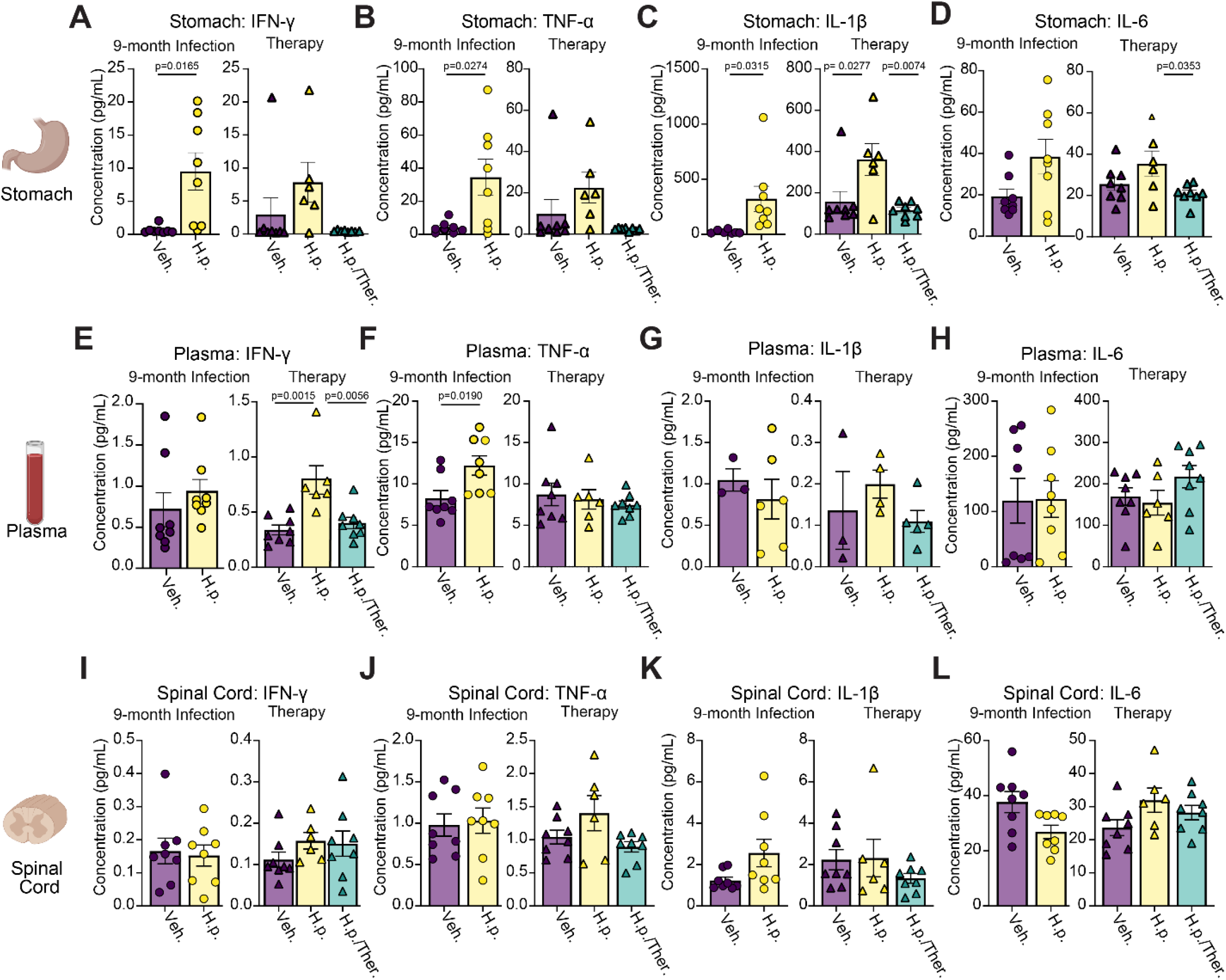
*H. pylori* infection induces local inflammation and systemic elevation of IFN-γ levels. Multiplexed cytokine assay showing the calculated concentration of IFN-γ, TNF-α, IL-1ß, and IL-6 levels measured in the **A.-D**. stomach tissue, **E.-H**. blood plasma, and **I.-L**. spinal cord of 9-month Infection and Therapy cohorts. For the 9-month infection cohort, n = 8 (Veh.) or n = 8 (H.p.), Welch’s T-tests were performed for **A, B, C, D**, and **K**. Unpaired T-tests were performed for the remaining panels. For the Therapy cohort, n = 8 (Veh.), 6 (*H*.*p*.), or 8 (*H*.*p*./Ther.), one-way ANOVA with Tukey’s multiple comparisons test was performed. Individual mice with cytokine levels below assay detection limits were excluded from analysis. Statistical significance (p < 0.05) is indicated above relevant comparisons; all other differences are non-significant.

### *H. pylori* infection induces mild neuroinflammation and neurodegeneration of dopaminergic neurons in the substantia nigra

Previous studies have demonstrated the ability for IFN-γ to facilitate dopaminergic neuron death in response to MPTP or rotenone^36^. Removal of either IFN-γ or microglia in these models resulted in attenuated cell death, suggesting an important role for IFN-γ-induced dopaminergic neuron death. We therefore sought to understand if there were inflammatory glial responses or dopaminergic neuron death in the substantia nigra of mice infected with *H. pylori*. We stained 9-month Infection and therapy cohorts with astrocytic marker glial fibrillary acidic protein (GFAP) and microglial marker, Ionized Calcium Binding Adaptor Molecule 1 (Iba1, **Fig. 3A**). *H. pylori*-infected mice showed a significant elevation in the GFAP-positive area (**Fig. 3B**), but there was no change in the microglial size or count (**Fig. 3C**). Interestingly, GFAP area remained increased even in mice that had received triple therapy and cleared the *H. pylori* infection (**Fig. 3D, 3E**), while microglia were also not changed in this cohort (**Fig. 3F**).

**Figure 3.**
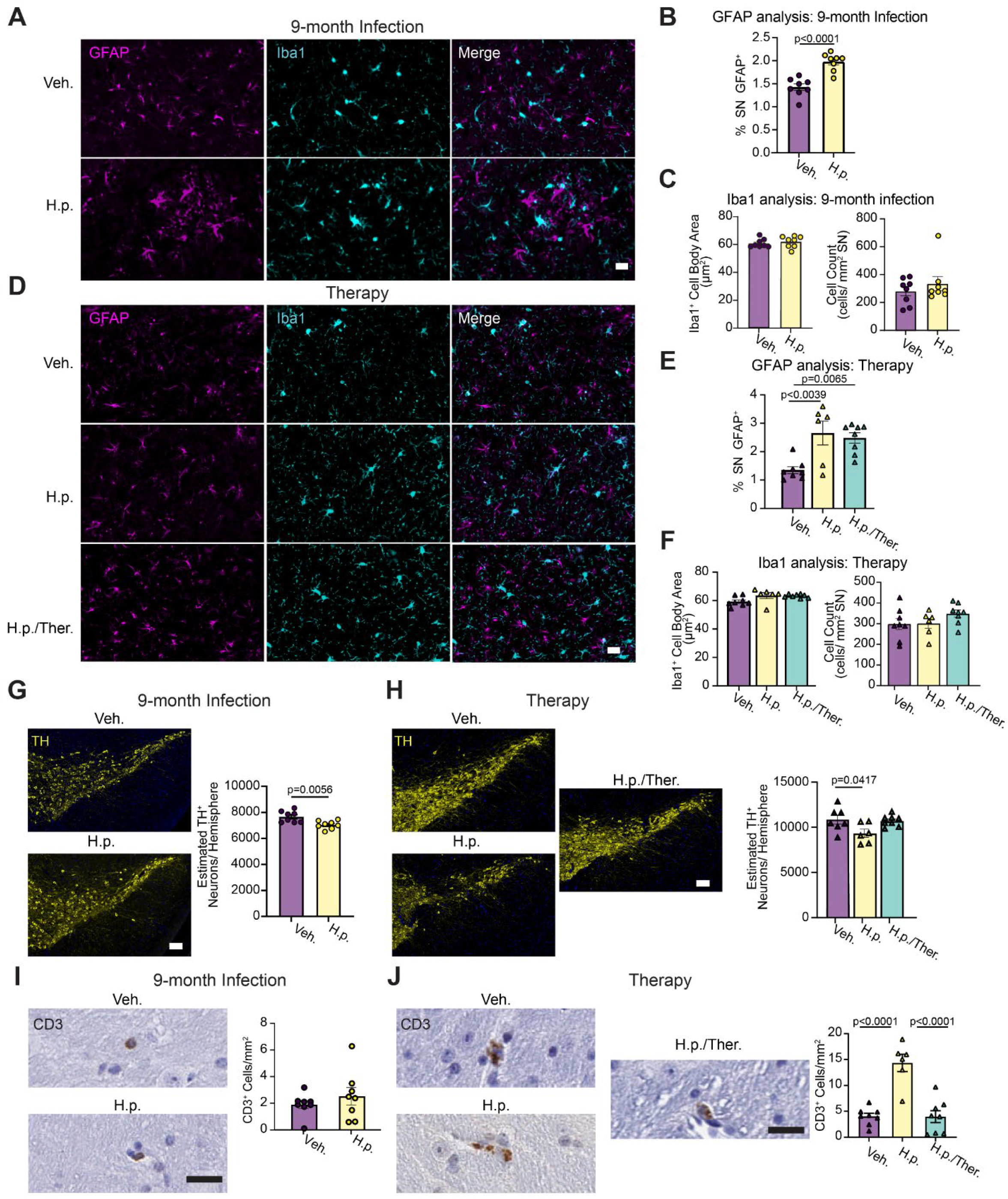
*H. pylori* infection leads to astrocytic activation and mild neurodegeneration in the substantia nigra. **A**. Representative images showing vehicle and *H. pylori* substantia nigra from the 9-month Infection cohort and stained for astrocytic marker GFAP and macrophage marker Iba1. Scale bar = 20 µm. **B**. Quantification of GFAP cell activation determined as % area of substantia nigra occupied in the 9-month Infection cohort. **C**. Quantification of microglial average Iba1^+^ cell body area and normalized Iba1^+^ cell count. **D**. Representative images showing vehicle and *H. pylori* substantia nigra stained for GFAP and Iba1 from the Therapy cohort. Scale bar = 20 µm. **E**. Quantification of GFAP signal measured as % area of substantia nigra occupied in the Therapy cohort. **F**. Quantification of microglial size and abundance. **G**. Representative images of 9-month Infection cohort substantia nigra stained for dopaminergic neuron marker tyrosine hydroxylase (TH), with quantification of estimated TH^+^ neurons per hemisphere. Scale bar = 100 µm. **H**. Representative images of Therapy cohort substantia nigra stained for TH with quantification of estimated TH^+^ neurons per hemisphere. Scale bar = 100 µm. **I**. Representative image of 9-month Infection cohort substantia nigra stained for CD3 with quantification of CD3^+^ cells plotted as cells per mm^2^ of substantia nigra. Scale bar = 20 µm. **J**. Representative image of Therapy cohort substantia nigra stained for CD3 with quantification of CD3^+^ cells plotted as cells per mm^2^ of substantia nigra. Scale bar = 20 µm. For the 9-month infection cohort, n = 8 (Veh.) or n = 8 (H.p.). A Welch’s T-test was performed for **I**. Unpaired T-tests were performed for the remaining panels. For the Therapy cohort, n = 8 (Veh.), 6 (*H*.*p*.), or 8 (*H*.*p*./Ther.), one-way ANOVA with Tukey’s multiple comparisons tests were performed. Statistical significance (p < 0.05) is indicated above relevant comparisons; all other differences are non-significant.

We next sought to understand the impact of *H. pylori* infection status on DA neurons in the substantia nigra. Sections throughout the substantia nigra were stained for tyrosine hydroxylase (TH) and neurons were counted to estimate the total number of neurons present. We observed a modest reduction in TH-positive neurons detected in *H. pylori-*infected mice after 9 months of infection (**Fig. 3G**). We observed a similar reduction of TH-positive neurons in *H. pylori*-infected mice in the therapy cohort, and triple therapy prevented this loss (**Fig. 3H**). Since we demonstrated that *H. pylori* infected mice do not develop α-synucleinopathy, this neuron loss is independent of α-synuclein pathology. IFN-γ is known to activate cytotoxic T-cells^38^, and T-cells have been documented to infiltrate the substantia nigra in PD^39-44^. Given the increased IFN-γ in *H. pylori-*infected mice, we hypothesized that T-cell infiltration could explain the reduction in DA neurons seen in *H. pylori*-infected mice. We stained the substantia nigra of vehicle and *H. pylori*-infected mice for CD3, a broad T-cell marker. Several T-cells were observed in the substantia nigra of mice (**Fig. 3I**). While there was no difference in the 9-month *H. pylori* cohort, there was a significant elevation in infiltrating T-cells in the *H. pylori*-infected animals in the therapy cohort, which was reversed by triple therapy (**Fig. 3J**). T-cells were sparse in both cohorts, and it is possible that the affected area was missed in sectioning for the 9-month cohort.

### *H. pylori* infection in a dual hit model

The odds ratio for *H. pylori* in relation to PD is 1.6^25^. The small amount of dopaminergic degeneration induced by *H. pylori* infection alone in our mouse model could be sufficient to confer an increased risk. However, we wanted to test an alternative hypothesis—*H. pylori* infection acts in concert with α-synuclein pathology thereby increasing risk only in those individuals with preexisting α-synucleinopathy. To test this hypothesis, we combined the *H. pylori* infection previously described with a seeded α-synucleinopathy model (**Fig. S4A**). As before, mice were treated with *H. pylori* or vehicle control at 3 months of age. One month after infection, mice were injected in the dorsal striatum with α-synuclein PFFs. The α-synuclein PFF model rapidly develops α-synuclein inclusions, followed by dopaminergic neuron death from 3-6 months post-injection (MPI)^45,46^. To capture pathology dynamics, mice were aged either 3 MPI or 6 MPI. Consistent with our previous results, *H. pylori* infection was sustained for the duration of the study (**Fig. 4B**) and induced a sustained infiltration of CD3^+^ T-cells in the mucosal layer of *H. pylori*-infected mice (**Fig. S4**). Consistent with our previous single-hit experiments, we found significant elevation in *H. pylori*-infected mice of several cyto- and chemokines, including IFN-γ, TNF-α, IL-1β, and IL-6, IL-2, IL-4, IL-5, IL-10, IL-12p70, and KC/GRO in the stomach of *H. pylori*-infect mice, although IL-6, IL-4, IL-5, and IL12p70 showed some difference between 3 and 6 MPI cohorts (**Fig. 4C-4F, S5A-S5F**). IFN-γ was also elevated in the plasma but not the spinal cord of *H. pylori*-infected mice (**Fig. 4G-4N, S5G-S5R**). No changes were observed in cytokine gene expression in the spinal cord either (**Fig. S6**).

**Figure 4.**
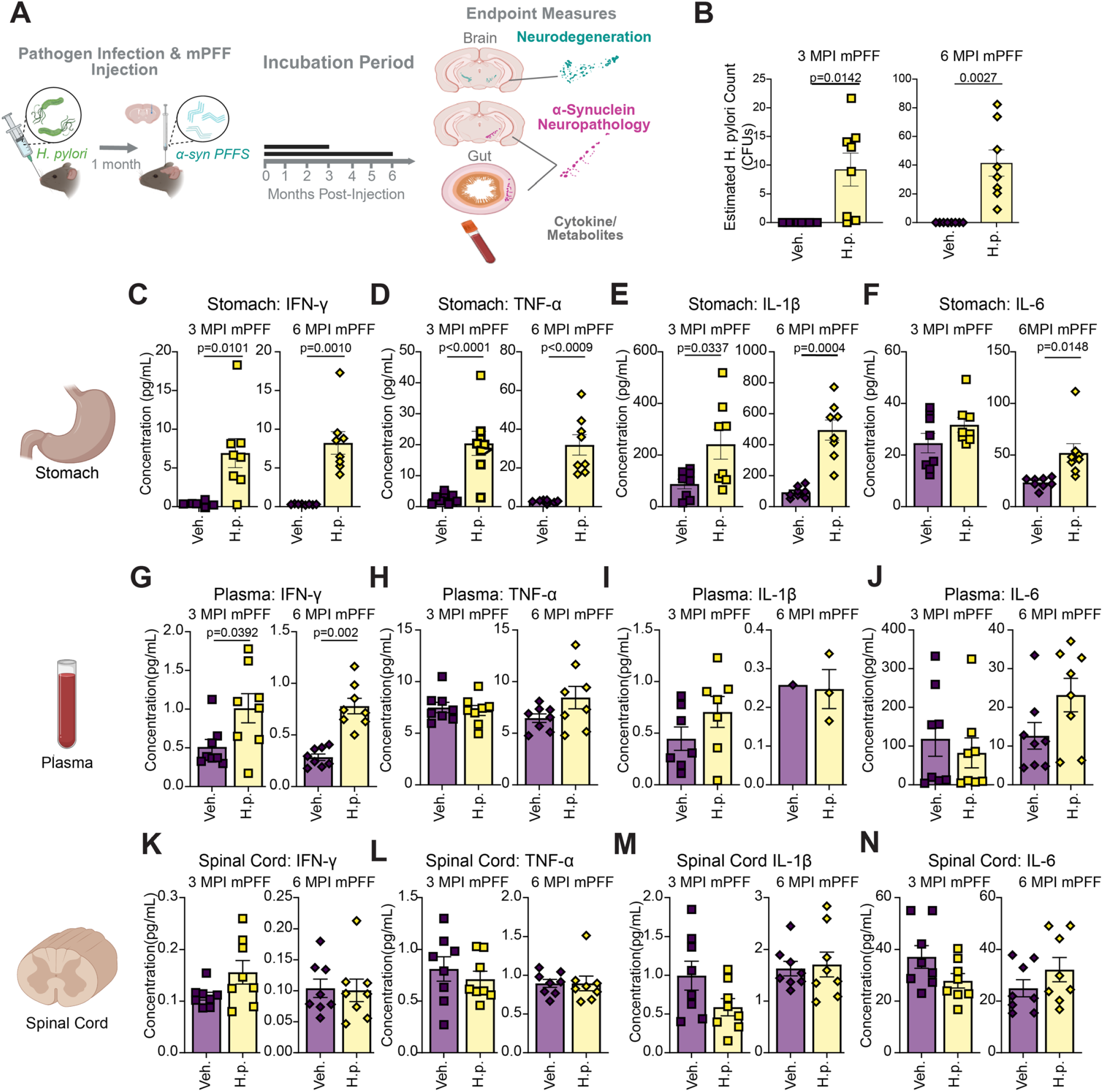
Local inflammation and elevated systemic INF-γ are also present in a dual-hit model of α-synuclein pathology. **A**. Experimental schematic. Mice were infected with *H. pylori* or fed the DMEM vehicle and injected in the dorsal striatum with mouse α-synuclein PFFs (PFF). Mice were subsequently aged 3 months or 6 months post mPFF injection. **B**. qPCR quantification of *H. pylori* load in 3 MPI mPFF cohort and 6 MPI mPFF cohorts. **C-N**. Multiplexed cytokine assay showing calculated concentration of IFN-γ, TNF-α, IL-1ß, and IL-6 levels measured in the **C.-F**. stomach tissue, **G.-J**. blood plasma, and **K.-N**. spinal cord tissue of the 3 MPI PFF and 6 MPI PFF cohorts. n = 8 (Veh.) or 8 (*H*.*p*.). Individual mice with cytokine levels below assay detection limits were excluded from analysis. A Welch’s T-test was performed for the **B, C, D, E, F, G, I**, and **K**. Unpaired T-tests were performed for all other panels. Statistical significance (p < 0.05) is indicated above relevant comparisons; all other differences are non-significant.

### *H. pylori* infection does not exacerbate α- synucleinopathy

To investigate whether *H. pylori* infection exacerbates the spread of α-synucleinopathy, we performed quantitative pathology analysis across the rostro-caudal axis of the mouse brain in a total of 1132 regions. All tissues were stained for pS129 α-synuclein and differences were evaluated between treatment cohorts. In both control and *H. pylori* infected mice, α-synuclein PFFs induce a broad distribution of pathology across the brain (**Fig. 5A, 5B**), primarily hitting regions that directly connect to the dorsal striatum early (cortex, amygdala, substantia nigra). The dorsal striatum itself is slower to develop pathology.

**Figure 5.**
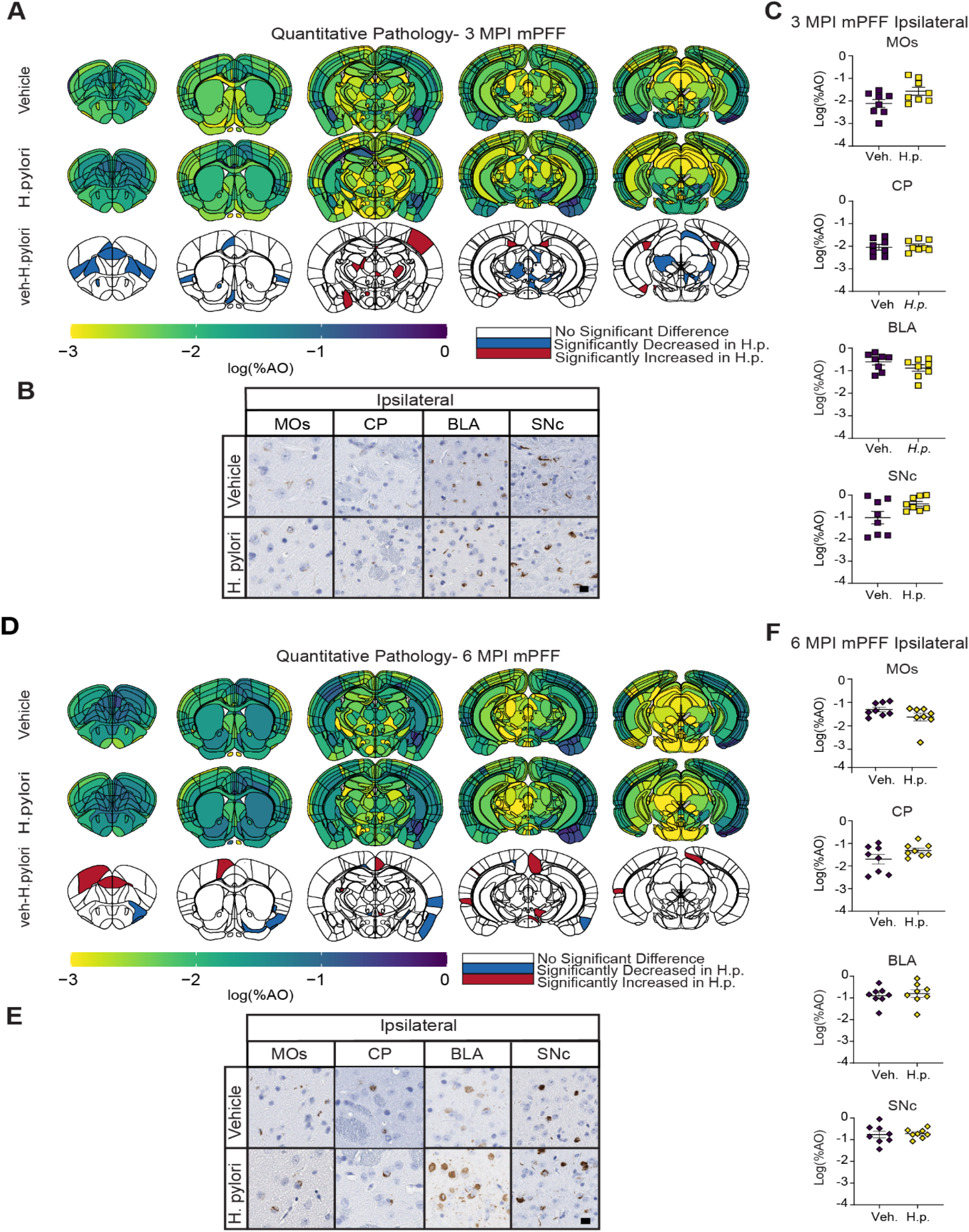
*H. pylori* infection does not meaningfully affect α-synuclein spread in a PFF injection model. **A**. Anatomic heatmaps of the mean regional α-synuclein pathology shown as log (% area occupied) at 3 MPI. Second-generation p-values calculated using ranged robust linear regression, of regional statistical significance of vehicle-treated mice compared to *H. pylori*-infected mice (pδ= 0). **B**. Representative images of selected regional α-synuclein pathology at 3 MPI. Scale bar = 10 µm. **C**. Quantification of selected brains regions. Regional α-synuclein shown as log (% area occupied). **D**. Anatomic heatmaps of the mean regional α-synuclein pathology shown as log (% area occupied) at 6 MPI. Second-generation p-values calculated using ranged robust linear regression, of regional statistical significance of vehicle-treated mice compared to *H. pylori*-infected mice (pδ = 0). **E**. Representative images of selected regional α-synuclein pathology at 6 MP. Scale bar = 10µm. **F**. Quantification of α-synuclein in selected brain regions. n= 8 (Veh.) or 8 (*H*.*p*.).

Overall, the pathology pattern observed in 3 MPI animals is consistent with previous studies^45,47^, and *H. pylori* infected mice showed no major deviations from the pathology pattern observed in vehicle treated mice (**Fig. 5C**). Pathology was more extensively distributed at 6 MPI (**Fig. 5D**), but again, no major differences were observed between treatment groups (**Fig. 5E, 5F**). These results suggest that *H. pylori* does not act synergistically to accelerate α-synucleinopathy.

### Neuroinflammation and neurodegeneration in *H. pylori*-infected, PFF-injected mice

To determine whether *H. pylori* together with α-synuclein pathology led to increased glial inflammation in the dual-hit model, we stained the brains of 3 MPI α-synuclein cohorts for Iba1 and GFAP (**Fig. 6A**). GFAP percent area occupied in the substantia nigra was increased in *H. pylori*-infected mice (**Fig. 6B**), similar to what we previously observed in mice without α-synuclein PFF injection. Additionally, there was a slight but significant increase in the Iba1+ cell body area in the substantia nigra of 3 MPI *H. pylori*-infected mice, (**Fig. 6C**). These data suggest that *H. pylori* is the primary driver of inflammation in this region and that α-synuclein pathology does not substantially change this pattern. Even at 6 MPI, when the substantia nigra has begun degenerating, there is still a similar difference between vehicle and *H. pylori* treated mice (**Fig. 6D-6F**).

**Figure 6.**
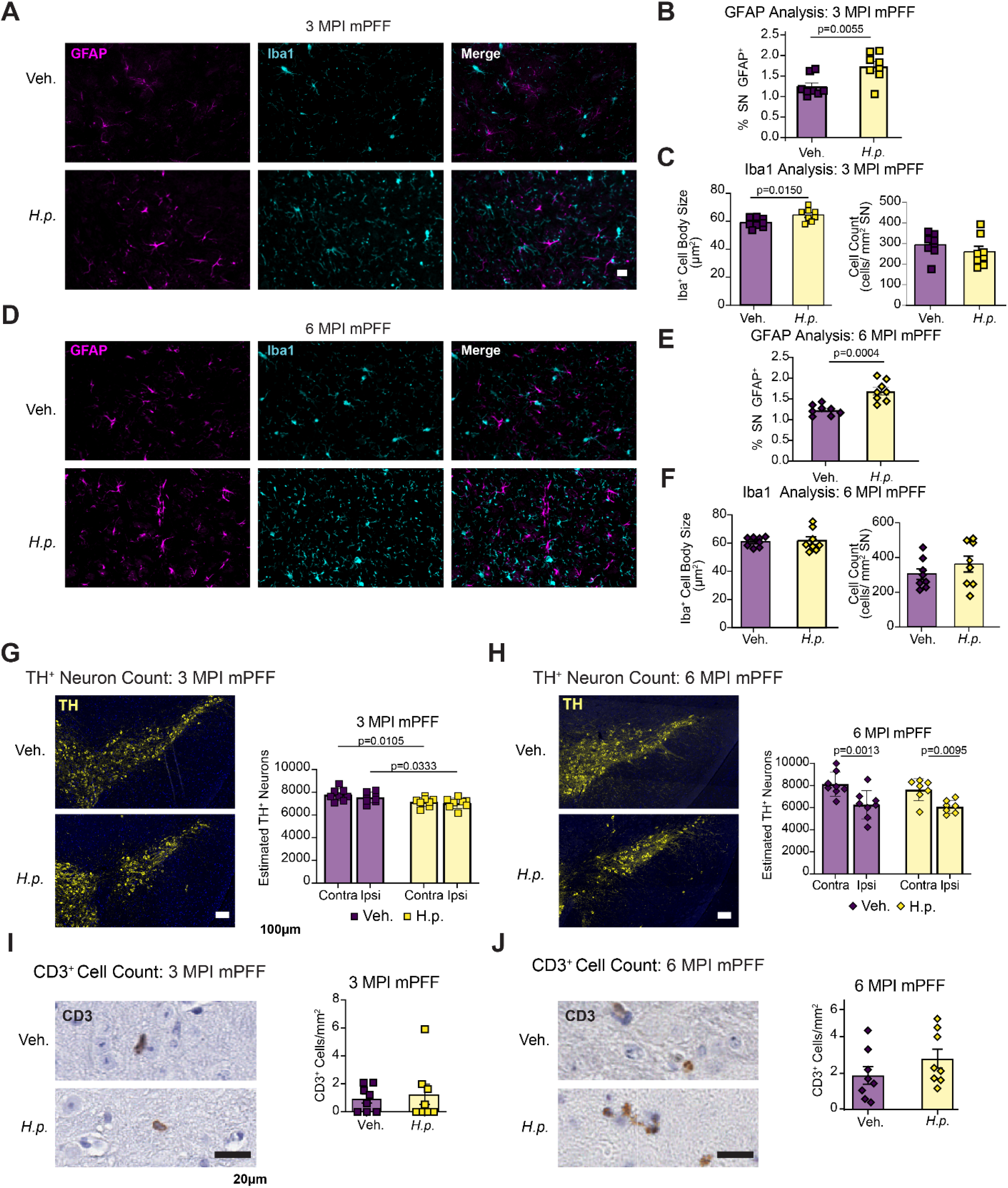
Neuroinflammation and neurodegeneration in *H. pylori*-infected, PFF-injected mice. **A**. Representative images showing vehicle and *H. pylori* substantia nigra from the 3 MPI PFF cohort stained for astrocytic marker, GFAP, and macrophage marker Iba1. Scale bar = 20 µm **B**. Quantification of GFAP % area of substantia nigra occupied in 3 MPI PFF cohort. **C**. Quantification of average Iba1^+^ cell body area and normalized Iba1^+^ cell count. **D**. Representative images showing vehicle and *H. pylori* substantia nigra stained for GFAP and Iba1 from the 6 MPI PFF mice. Scale bar = 20 µm. **E**. Quantification of GFAP signal measured as % area of substantia nigra occupied in the 6 MPI mPFF cohort. **F**. Quantification of average Iba1^+^ cell body area and normalized Iba1^+^ cell count. **G**. Representative images of 3 MPI PFF cohort substantia nigra stained for tyrosine hydroxylase (TH), with quantification of estimated TH^+^ neurons separated into ipsilateral and contralateral hemispheres. Scale bar = 100 µm. **H**. Representative images of 6 MPI cohort substantia nigra stained for TH with quantification of estimated TH^+^ neurons separated into ipsilateral and contralateral hemispheres. Scale bar = 100 µm. **I**. Representative image of 3 MPI PFF substantia nigra stained for CD3 with quantification of CD3^+^ cells plotted as cells per mm^2^ of substantia nigra. Scale bar = 20µm. **J**. Representative image of 6 MPI PFF cohort substantia nigra stained for CD3 with quantification of CD3^+^ cells plotted as cells per mm^2^ of substantia nigra. Scale bar = 20 µm. n = 8 (Veh.) or 8 (*H*.*p*.), A Welch’s T-test was performed **F**. and **I**. Two-way ANOVA with uncorrected Fisher’s LSD test was performed for **G**. and **H**. Unpaired T-tests were performed for all other panels. Statistical significance (p < 0.05) is indicated above relevant comparisons; all other differences are non-significant.

To assess the impact of two hits on neuron survival, we quantified dopaminergic neurons through the substantia nigra. Similar to the *H. pylori* alone cohorts, there was a significant loss of neurons on both sides of the brain of *H. pylori* infected mice at 3 MPI (**Fig. 6G**). There was no significant α-synuclein-induced loss at this timepoint, which would appear as a loss of neurons ipsilateral to the injection site. By 6 MPI, there was a substantial loss of DA neurons ipsilateral to the α-synuclein PFF injection site, but this loss was similar between vehicle and *H. pylori* treated animals (**Fig. 6H**), suggesting again that there is not a synergistic effect between *H. pylori* infection and α-synuclein pathology.

Finally, we assessed CD3^+^ T-cell infiltration into the substantia nigra. Like in the 9-month Infection and therapy mice, few CD3^+^ T-cells were present, and there was no significant difference between vehicle and *H. pylori* infected mice (**Fig. 6J**).

## DISCUSSION

There is significant epidemiological evidence supporting a link between *H. pylori* infection and PD pathogenesis. However, the link between *H. pylori* infection and the development and progression of α-synucleinopathy has yet to be investigated in animal models. In this study, we sought to understand the contributions of *H. pylori* to PD-relevant phenotypes. We successfully established a chronic *H. pylori* infection that was cleared following antibiotic triple therapy. Our findings suggest that *H. pylori* infection alone is insufficient to trigger the formation of α-synuclein pathology within 9 months. Further, we found that *H. pylori* infection does not change the initiation or progression of α-synucleinopathy in a seed-based model at the timepoints tested. However, we found that *H. pylori* infection does result in chronic localized inflammation that extends to the plasma and can lead to astrocytic activation and T-cell infiltration in the substantia nigra, and mild neurodegeneration of dopaminergic neurons. These findings support a role for infectious diseases, such as *H. pylori*, in inflammation-driven neurodegeneration, which may occur independently or in parallel with proteinopathies.

We observed local elevations, especially in IFN-γ, TNF-α, and IL-1β in the stomach and an increase in plasma IFN-γ levels in *H. pylori*-infected mice with and without PFF injections. The elevation in IFN-γ we observed is consistent with reports of PD patients presenting with increased IFN-γ in plasma^36^. Mice deficient in IFN-γ showed attenuated dopaminergic neuron loss following MPTP treatment, suggesting an important role of IFN-γ in neuronal toxicity. Further, IFN-γ can damage dopaminergic neurons in culture, but only when co-cultured with microglia. Additionally, KC/GRO (also known as CXCL1) is induced by gut microbiota in mice and triggers neuroinflammation and deposition of iron in the substantia nigra leading to neuronal death^48^. Finally, IL-2 is a T-cell activating cytokine^49^ and is seen to be elevated in *H. pylori* infected mice. Together, these results indicate *H. pylori* is responsible for the induction of a chronic inflammatory phenotype that spreads from the stomach throughout the body and may help create conditions favorable to PD pathogenesis.

Astrocytes play a prominent role in neuroinflammation and maintenance of the blood-brain barrier^50^. Astrocytes can become reactive during pathological conditions. Reactive astrocytes take on a neurotoxic phenotype, exhibit increased expression of GFAP, and release proinflammatory cytokines, contributing to neuroinflammation^51^. The astrocytic activation and neuronal damage we observed in *H. pylori* mice could be driven by circulating cytokines, but another report also suggested that outer membrane vesicles secreted by *H. pylori* can cross the blood-brain barrier, trigger astrocyte activation through NF-κB-dependent pathways and facilitate neuronal damage^52^.

We also identified increased CD3^+^ T-cells in the substantia nigra and brain parenchyma of some, but not all, *H. pylori* infected cohorts. T-cells have been reported to play a role in PD pathogenesis. CD4^+^ and CD8^+^ T-cells infiltrate the substantia nigra in rats injected with AAV-α-synuclein, leading to microglial upregulation of MHC-II correlating with dopaminergic neuron death and resulting in motor deficits^53^. Neurons in the substantia nigra express MHC-I in response to IFN-γ secreted by microglia resulting in neuron death mediated by cytotoxic T cells^54^. Additional compelling studies have shown that circulating T-cells in PD can recognize α-synuclein epitopes, resulting in T-cell activation^55^. These results suggest that PD could involve an auto-immune component.

There is historical precedent for the hypothesis that infection could independently drive loss of dopaminergic neurons. There are several historical cases of parkinsonism arising directly as a result of infectious disease. Notably, the “Spanish flu” pandemic of the early 1900s was linked to an outbreak of encephalitis lethargica^56^. Patients who survived developed post-encephalitic parkinsonism that encompassed many of the cardinal features of PD, albeit with more severe symptoms that could include catalepsy. These patients could also be “awakened” by levodopa, suggesting that this disease was related to dopaminergic neuron dysfunction or death^57,58^. This long history of violent encephalitis’ induction of parkinsonism could hint towards a particular vulnerability of the mesencephalon and dopaminergic neurons in general that underlies the potential for infectious diseases to increase the propensity for PD. This hypothesis was tested at an unprecedented scale during the COVID-19 pandemic. Case reports from early in the pandemic suggested the emergence of post-COVID parkinsonism^59-61^. Despite these early suggestions, COVID-19 has not been conclusively linked to development of PD^61-64^, though longer periods may be required between an initial infectious insult and symptom emergence. It is important to note that COVID-19 is not thought to induce encephalopathy^63^, and even post-encephalitic syndromes are largely non-progressive, which separate them from PD. However, if there was a sub-threshold assault on the dopaminergic system, coupled with aging process, it could lead to increased incidence of developing PD.

The current study has several limitations. Importantly, mice have a limited lifespan, so our chronic infection with *H. pylori* (9 months) would be a relatively short time period for humans. It is therefore possible that the small phenotypes observed in mice could be exacerbated in individuals that sustain years-long infections. Second, we used one inbred mouse strain. Humans are much more genetically diverse and there have been studies associating specific *Il1b* genotypes with increased susceptibility to the pathogenic effects of *H. pylori*^65-69^. It may be the case that the mouse strain used in this study is particularly resistant to pathogenic effects of *H. pylori*.

Our study identifies one means by which infectious diseases, such as *H. pylori* infection, can lead to degeneration of vulnerable neurons, thereby increasing the lifetime risk of developing PD. Given the previous associations of infectious disease and neuroinflammation with PD^70,71^, it is interesting to consider whether there is a specific subset of PD cases that have an inflammatory etiology^72^ and may be more responsive to anti-inflammatory or eradication therapies. There is an epidemiological association between *H. pylori* eradication and decreased PD severity^73-75^. How would we identify such a subset of inflammation-related PD? In living cohorts, serology could be used determine whether certain people have elevated cytokines^76^. Notably, in the studies which have reported T-cell infiltration into the brains of PD patients, there is never strong specificity of this profile, with many PD patients showing similar T-cell levels in the brain to unaffected controls. It would be interesting to perform serology on these same cases to understand if there is a specific infectious or inflammatory marker present in those cases that do exhibit T-cell infiltration.

Our findings reinforce the significance of the gut-brain axis in PD pathogenesis, where gut microbiota and systemic inflammation may create a permissive environment for neurodegeneration. Further understanding the interaction between infectious agents, the immune system, and the central nervous system could facilitate the discovery of novel therapeutic strategies and improve outcomes for PD patients.

## MATERIALS AND METHODS

### Animals

All housing, breeding, and procedures were performed according to the NIH Guide for the Care and Use of Experimental Animals and approved by the Van Andel Institute Institutional Animal Care and Use Committee. Both male (n=35) and female (n=35) mice were used and were 3 months old at the time of *H. pylori infection* and 4 months old at the time of PFF injection or antibiotic triple therapy treatment. No influence of sex was identified in the measures reported in this study.

### *H. pylori* culture, infection, and antibiotic triple therapy treatment

The *H. pylori* Pre-Mouse Sydney Strain 1 (PMSS1) strain bacteria were a gift from Dr. Manuel Amieva (Stanford). *H. pylori*^27,28^ PMSS1 was cultured from frozen glycerol stock onto horse blood agar plates and incubated in a Campy jar with Campy Pak (sachet) at 37°C (Ex. Jar--GasPak™ 100 Anaerobic System Item # 801340; sachet--BD EZ Campy container system #260680) for 2 days. One day before infection, the bacteria were suspended in a 3 mL solution of Brucella broth with 10% fetal bovine serum (FBS). The suspension was incubated in an incubator at 37°C for at least 2 hours. Bacteria were quantified by OD600 measurement and diluted to an OD600 of 0.018 in 11 mL of fresh Brucella broth with 10% FBS. This second culture suspension was incubated under the same conditions overnight for 17-18 hours. Cultured bacteria were quantified by OD600 measurement and resuspended to a concentration of 1X10^8^ CFU/5 μL in co-culture media (DMEM +5% FBS + 10% Brucella broth). Mice were fed 5 μL of the suspension in DMEM (1X10^8^ CFUs total).

4-month-old mice were treated with triple antibiotic therapy 1 month after initial *H. pylori* infection. Mice received 20 mL/kg Omeprazole (36.5 mg/ml), Amoxicillin (23.33 mg/ml), and Clarithromycin (11.17 mg/ml) in PBS by oral gavage once per day for 5 days.

### α-synuclein PFFs

Purification of recombinant mouse α-synuclein and generation of α-synuclein PFFs was conducted as described elsewhere^77,78^. The pRK172 plasmid containing the gene of interest was transformed into BL21 (DE3) RIL-competent *E. coli* (Agilent Technologies Cat#230245). A single colony from this transformation was expanded in Terrific Broth (12 g/L of Bacto-tryptone, 24 g/L of yeast extract 4% (vol/vol) glycerol, 17 mM KH_2_PO_4_ and 72 mM K_2_HPO_4_) with ampicillin. Bacterial pellets from the growth were sonicated and the sample was boiled to precipitate undesired proteins. The supernatant was dialyzed with 10 mM Tris, pH 7.6, 50 mM NaCl, 1 mM EDTA, 1 mM phenylmethylsulphonyl fluoride (PMSF) overnight. Protein was filtered with a 0.22 µm filter and concentrated using Vivaspin 15R 10K centrifugal filters (Sartorius VS15RH01). Protein was then loaded onto a Superdex 200 column and 2 mL fractions were collected. Fractions were run on SDS-PAGE and stained with InstaBlue protein stain (Fisher Scientific 50-190-5499) to select fractions that were highly enriched in α-syn. These fractions were combined and dialyzed in 10 mM Tris, pH 7.6, 50 mM NaCl, 1 mM EDTA, 1 mM PMSF overnight. Dialyzed fractions were applied to the MonoQ column (Cytiva, HiTrap Q HP 17115401) and run using a linear gradient from 25 mM NaCl to 1 M NaCl. Collected fractions were run on SDS-PAGE and stained with InstaBlue protein stain. Fractions that were highly enriched in α-synuclein were collected and dialyzed into DPBS. Protein was filtered through a 0.22 µm filter and concentrated to 7 mg/mL (α-syn) with Vivaspin 15R 10K centrifugal filters. Monomer was aliquoted and frozen at -80°C. For preparation of α-synuclein PFFs, α-synuclein monomer was shaken at 1,000 rpm for 7 days. Conversion to PFFs was validated by sedimentation at 100,000 x *g* for 60 minutes and by thioflavin S staining.

### α-synuclein PFF stereotaxic injections

α-Synuclein PFFs were vortexed and diluted with DPBS to 2 mg/mL. α-Synuclein PFFs were sonicated in a cooled bath sonicator at 9°C (Diagenode Bioruptor® Pico; 10 cycles; setting medium; 30 seconds on, 30 seconds off). Mice were injected at 4 months of age. Mice were deeply anesthetized with isoflurane and injected unilaterally by inserting a single needle into brain targeting the right dorsal striatum (coordinates: +0.2 mm relative to Bregma, +2.0 mm from midline, 2.5 mm beneath the dura) with 5 µg α-synuclein PFFs (2.5 µL). Injections were performed using a 10 µL syringe (Hamilton 7635-01, NV) with a 34-gauge needle (Hamilton 207434, NV) at a rate of 0.4 µL/minute.

### Multiplexed cytokine analysis

Secretion of inflammatory cytokines Interferon gamma (IFN-γ), Interleukin (IL)-1β, IL-2, IL-4, IL-5, IL-6, IL-10, IL-12, tumor necrosis factor alpha (TNF-α), and chemokine keratinocyte chemoattractant (KC/GRO) were assessed for all experimental and control mice using the MSD® V-PLEX Proinflammatory Panel 1 Mouse Kit (Catalog # K15056D-2). Stomach and spinal cord tissues, each weighing between 30 and 40 mg were homogenized in RIPA lysis buffer(50mM Tris-HCl, pH 7.6, 150 mM NaCl, 10% NP-40, 0.5% sodium deoxycholate, 0.1% SDS,1 mM EDTA, pH 8.0, 10mM NaF) at a ratio of 6 times the tissue weight (v/w) using a Dounce hand homogenizer. The lysis buffer was supplemented with 5 mM sodium orthovanadate, 5 mM PMSF, 5 mM phosphatase inhibitor cocktail 3 (Sigma Aldrich P0044), and approximately ½ tablet of Pierce protease inhibitor mini tablets (Thermofisher A32955). Samples were sonicated in the Diogenode Biorupter Pico bath sonicator for 30 seconds on and 30 seconds off for 10 cycles. According to the manufacturer’s recommendation, plasma was thawed and diluted in MSD diluent 41 solution. 50 µL of each tissue homogenate or plasma sample was diluted in 150 µL of diluent 41 to create a 200 µL sample solution with a 1:4 dilution ratio.

The provided multispot plate was initially washed 3 times with 150 µL/well of wash buffer (PBS + 0.05% Tween-20). 50 µL/well of prepared samples, calibrator standards, or controls was added. The plate was sealed with an adhesive plate seal and incubated at room temperature for 2 hours while shaking at 700 rpm. The plate was washed 3 more times with 150 µL/well of wash buffer. 25 µL of detection antibody was added to each well. The plate was sealed and incubated at room temperature for 2 hours while shaking at 700 rpm The plate was washed 3 final times with 150 µL/well of wash buffer. Finally, Read Buffer T was added to each well and the plate was immediately read in the MSD QuickPlex SQ 120 instrument.

The median lower limits of detection (LLOD) for the MSD V-PLEX assay were as follows: IFN-γ, 0.04 pg/mL; IL-1β, 0.11 pg/mL; IL-2, 0.22 pg/mL; IL-4, 0.11 pg/mL; IL-5, 0.06 pg/mL; IL-6, 0.61 pg/mL; IL-10, 0.94 pg/mL; IL-12p70, 9.95 pg/mL; KC/GRO, 0.24 pg/mL; and TNF-α, 0.13 pg/mL. In stomach tissue, all samples were within detection limits except for one sample in the IL-12p70 assay. In plasma, the number of samples below detection limits were as follows: IL-1β (n = 32), IL-2 (n = 43), IL-4 (n = 43), IL-6 (n = 17), and IL-12p70 (n = 46). For spinal cord tissue, one sample was below detection limits for IFN-γ, IL-1β, IL-5, IL-6, IL-10, IL-12p70, KC/GRO, and TNF-α; two samples were below detection limits for IL-2; and three samples were below detection limits for IL-4. Samples with cytokine concentrations below the LLOD were excluded from statistical analyses.

### QPCR

*H. pylori* bacterial load was assessed by qPCR of the *H. pylori* gene *glmM*. DNA from stomach samples was extracted using the QIAamp® DNA mini kit (Qiagen 51304). A reaction mixture was prepared at a final volume of 20 µL containing 10 µL of PowerTrack™ SYBR Green master mix kit (ThermoFisher Scientific A46109), 0.5µL of yellow sample buffer (ThermoFisher Scientific A46109), 0.2µM *glmM* forward primer, 0.2µM *glmM* reverse primer, 2 ng template DNA (2µL total volume), and RNase-free water. Mouse *Gapdh* was used as a reference gene. The reaction was performed on the QuantStudio™ 3 Real-Time PCR System (Thermofisher Scientific). PCR amplification consisted of a preamplification step and a relative quantification step. The preamplification reaction was performed with an initial denaturation step at 95°C for 3 minutes followed by 16 amplification cycles at 94°C for 1 min, 55°C for 1 min, and 72°C for 2 min. The relative quantification reaction was performed with an initial denaturation step at 95°C for 3 min followed by 40 cycles at 95°C for 10 seconds, 60°C for 40 seconds, and 95°C for 15 seconds. The results of each reaction were validated by visualization on a DNA agarose gel.

Proinflammatory cytokine gene expression was monitored in spinal cord tissues by qPCR. IL-1 β *(Il1b)*, IL-2 (*Il2)*, IL-6 (*Il6)*, IL-17 (*Il17a)*, IFN-γ *(Ifng)*, and TNF-α *(Tnf)* were assessed. RNA from spinal cord samples was extracted with the Qiagen RNeasy® mini kit (Qiagen 74106). A reaction mixture was prepared with the SuperScript™ III One-Step RT-PCR System with Platinum™ Taq DNA Polymerase system (ThermoFisher Scientific 12574026) at a final volume of 20 µL. The reaction mixture contained 0.4 µL SuperScript™ III RT/Platinum™ Taq Mix, 10 µL of 2x SYBR Green ™ reaction mix, 0.2 µL ROX ™ passive reference dye, 1 nM forward primer, 1 nM reverse primer, 1 ng RNA template (2µL total volume), and nuclease free water. β-actin (*Actb)* was used as a reference gene. The reaction was performed on the QuantStudio™ 3 Real-Time PCR System (Thermofisher Scientific). The PCR reaction consisted of an initial step at 50°C for 2 minutes followed by 95°C for 10 minutes and 40 cycles at 95°C for 15 seconds, 60°C for 1 minute. Relative quantities of each cytokine were calculated using the equation: Fold change = 2 ^-ΔΔCT^ where ΔCT = avgCt_cytokine_ -avgCT_β-actin_ and ΔΔCT = ΔCT_*H*.*pylori*_ -ΔCT_control_.

**Table 1.**
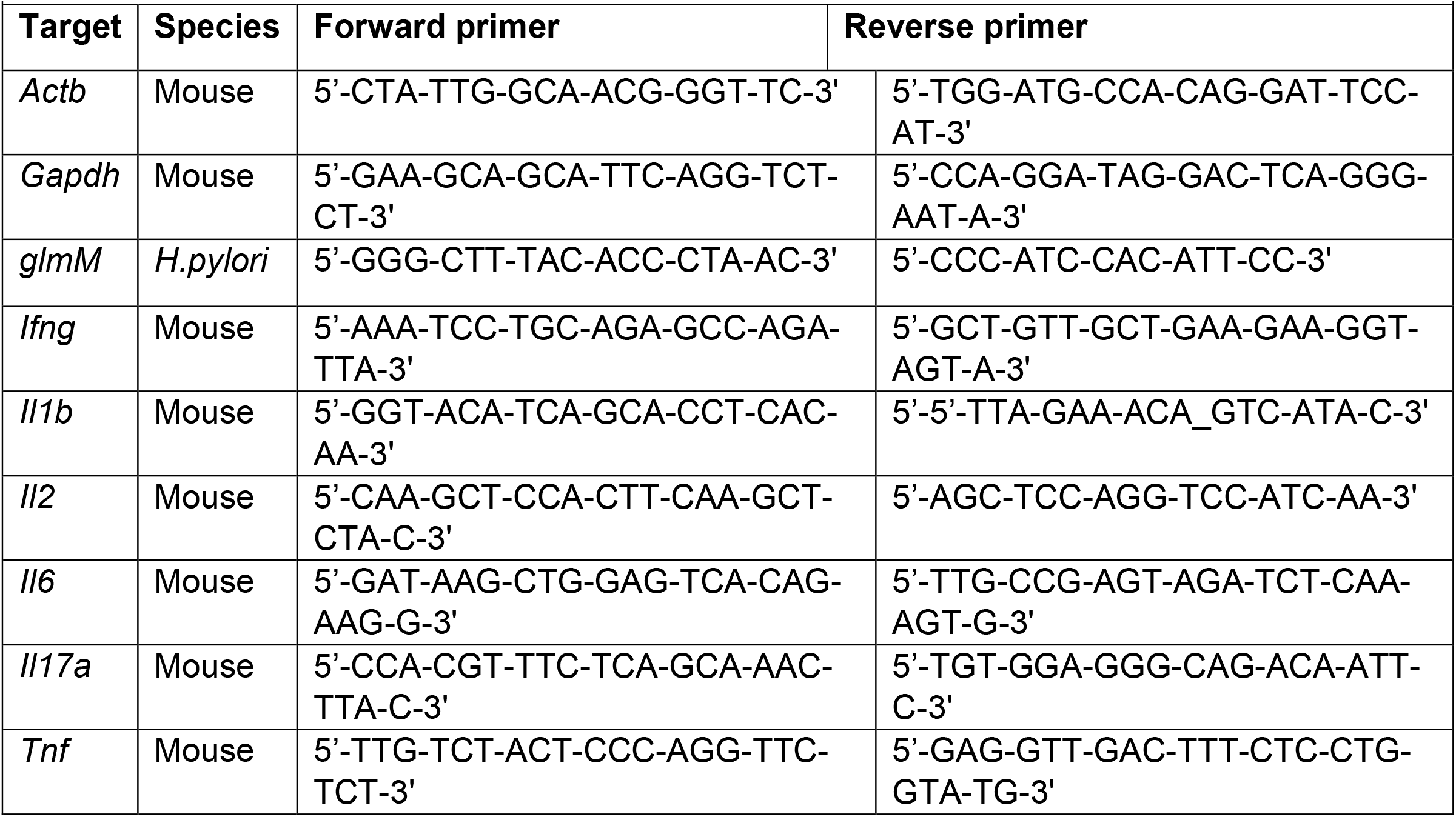
List of primers used.

### Tissue Collection

Mice were perfused transcardially with phosphate-buffered saline (PBS), and brains, stomach, spinal cords, and plasma were collected. The stomachs were opened by cutting along the lesser curvature and cut into left and right halves^79^. The left halves of the stomachs, spinal cords, and plasma were flash frozen in liquid nitrogen and stored at - 80°C. Brains and the right halves of stomach underwent overnight fixation in 4% paraformaldehyde (PFA). Brains and the right halves of the stomachs were processed into paraffin via sequential dehydration and perfusion with paraffin under vacuum (70% ethanol for 1 hour, 80% ethanol for 1 hour, 2 times 95% ethanol for 1 hour, 3 times 100% ethanol for 1 hour, 2 times xylene for 30 minutes, paraffin for 30 minutes at 60 °C, paraffin for 45 minutes at 60 °C). Brains were then embedded in paraffin blocks, cut into 6 µm sections and mounted on glass slides.

### Immunohistochemistry and immunofluorescence

Slides were de-paraffinized with 2 sequential 5-minute washes in xylenes, followed by 1-minute washes in a descending series of ethanol solutions: 100%, 100%, 95%, 80%, 70%. Slides were then incubated in deionized water for one minute and transferred to antigen unmasking solution (Vector Laboratories; Cat# H-3300) and microwaved for 15 minutes at 95°C (BioGenex EZ-Retriever System). Slides were allowed to cool for 20 minutes at room temperature and washed in running tap water for 10 minutes. To stain pS129 α-syn, slides were treated with proteinase K (10 µg/mL) for 5 minutes followed by 5 mM PMSF for an additional 5 minutes to quench proteinase activity and were washed in running tap water for 10 minutes. Slides were incubated in 7.5% hydrogen peroxide in water to quench endogenous peroxidase activity. This step was skipped for immunofluorescence. Slides were washed for 10 minutes in running tap water, 5 minutes in 0.1 M Tris (diluted from 0.5 M Tris made from tris base and concentrated hydrochloric acid to pH 7.6), then blocked in 0.1 M Tris/2% fetal bovine serum (FBS) for 1-2 hours. Slides were incubated in primary antibody in 0.1 M Tris/2% FBS in a humidified chamber overnight at 4°C.

For immunohistochemistry, the primary antibody was rinsed off with 0.1 M Tris and incubated in 0.1M Tris for 5 minutes, then incubated with goat anti-rabbit (Vector BA1000, RRID: AB_2313606) or horse anti-mouse (Vector BA2000, RRID: AB_2313581) biotinylated IgG in 0.1 M Tris/2% FBS 1:1000 for 1 hour. The biotinylated antibody was rinsed off with 0.1 M Tris for 5 minutes, then incubated with avidin-biotin solution (Vector PK-6100, RRID: AB_2336819) for 1 hour. Slides were then rinsed with 0.1M Tris and washed for 5 minutes with 0.1 M Tris. The slides were then developed with ImmPACT DAB peroxidase substrate (Vector SK-4105, RRID: AB_2336520) and counterstained briefly with Harris Hematoxylin (Fisher 67-650-01). Slides were washed in running tap water for 5 minutes, dehydrated in ascending ethanol for 1 minute each (70%, 80%, 95%, 100%, 100%), then washed twice in xylenes for 5 minutes. The slides were mounted using Cytoseal Mounting Media (Fisher, Cat# 23-244-256). Slides were scanned into digital format on an Aperio AT2 microscope using a 20x objective (0.75 NA) into ScanScope virtual slide (.svs) files.

For immunofluorescence, slides were incubated in secondary antibodies (see list of secondary antibodies below) in the dark for 2 hours at room temperature. Slides were washed 3 times for 10 minutes in 0.1 M Tris in the dark. Slides were mounted with coverslips in ProLong gold with DAPI (Invitrogen, Cat#P36931). Fluorescent slides were imaged at 20x magnification on a Zeiss AxioScan microscope.

**Table 2.** List of primary antibodies used.

| **Table2. List of primary antibodies used** |  |  |  |  |
| --- | --- | --- | --- | --- |
| **Antigen** | **Company** | **Catalog Number** | **Dilution** | **RRID** |
| CD3 epsilon | Abcam | Ab16669 | 1:1,000 | AB_443425 |
| Choline Acetyltransferase | Abcam | Ab178580 | 1:1,000 | AB_2721842 |
| GFAP (2.2B10) | ThermoFisher | 13-0300 | 1:1,000 | AB_2532994 |
| *H. pylori* | Abcam | Ab172611 | 1:1,000 | AB_2891050 |
| Iba1 | Fujifilm Wako | 019-19741 | 1:500 | AB_839504 |
| pS129 α-synuclein | Abcam | Ab51253 | 1:20,000 | AB_869973 |
| Tyrosine Hydroxylase | Sigma-Aldrich | T2928 | 1:5,000 | AB_477569 |

**Table 3.** List of secondary antibodies used.

| **Table 3. List of secondary antibodies used** |  |  |  |  |
| --- | --- | --- | --- | --- |
| **Antibody name** | **Company** | **Catalog Number** | **Dilution** | **RRID** |
| Horse anti-mouse IgG, biotinylated | Vector Laboratories | BA-2000 | 1:1,000 | AB_2687893 |
| Goat anti-rabbit IgG, biotinylated | Vector Laboratories | BA-1000 | 1:1,000 | AB_2916034 |
| Goat anti-mouse IgG1, Alexa Fluor 488 | ThermoFisher | A-21121 | 1:500 | AB_2535764 |
| Goat anti-mouse IgG (H+L), Alexa Fluor 546 | ThermoFisher | A-11003 | 1:500 | AB_2534071 |
| Goat anti-mouse IgG1, Alexa Fluor 647 | ThermoFisher | A-21240 | 1:500 | AB_2535809 |
| Goat anti-mouse IgG2a, Alexa Fluor 488 | Invitrogen | A21131 | 1:500 | AB_2535771 |
| Goat anti-rabbit IgG (H+L), Alexa Fluor 546 | Invitrogen | A11010 | 1:500 | AB_2534077 |
| Goat anti-rabbit IgG (H+L), Alexa Fluor 647 | ThermoFisher | A-21244 | 1:500 | AB_2535812 |
| Goat anti-rat IgG (H+L), Alexa Fluor 546 | Invitrogen | A11081 | 1:500 | AB_2534125 |

### Cell quantification

To assess dopaminergic cell loss in the substantia nigra, every 10^th^ slide through the midbrain was stained according to the immunofluorescence protocol described above for tyrosine hydroxylase. Section selection and image analysis were done blinded to treatment. Section images were imported into the open-access image analysis software, QuPath^80^ (RRID SCR_018257). Regions of interest comprising the left and right substantia nigra were annotated on each section. Cell detection was applied to detect DAPI-stained nuclei with an intensity threshold above 150 and a cell expansion of 5 µm. An automated object classifier was trained to identify detected cells with the tyrosine hydroxylase signal inside the cell body. The results of the automated object classifier were validated via manual counting.

To evaluate the activation of microglia and astrocytes in the substantia nigra, QuPath was used to measure Iba1 ^+^ cell body size and the percent area of the substantia nigra occupied by GFAP^+^ cells, respectively. Two midbrain sections 10 slides apart that contained the substantia nigra were selected and co-stained for Iba1 and GFAP. Iba1 cell body size was measured in QuPath with an automated cell detection set to detect Iba1 positive cells with fluorescence intensity threshold above 250 and a cell expansion of 0 µm. An automated object classifier was trained to distinguish between true positive and false positive detections. The average Iba1^+^ cell body size was calculated for each brain. For the GFAP activation analysis, a region of interest that contained the substantia nigra was annotated. The pixel classification feature of QuPath was used to create a thresholder that would identify GFAP^+^ cells with a fluorescent intensity above 1250 within the region of interest. For each brain, the total area of both regions of interest as well as the total area occupied by GFAP ^+^ cells were used to calculate the percentage of area occupied by GFAP staining.

To count CD3 cells in the substantia nigra and stomach tissue, a ROI was delineated around the substantia nigra and a 1 mm^2^ ROI was drawn around the mucosal layer of the stomach in QuPath. CD3-positive cells within the respective ROIs were counted manually using the points tool of QuPath.

### Pathology quantification

All section selection, registration, and quantification were done blinded to treatment. An adapted version of the QUINT workflow^81^ (RRID: SCR_023856) was used to quantify α-synuclein pathology and register brain sections to the Allen Brain Atlas CCFv3^82^. This workflow utilizes a suite of open-access programs for quantification and spatial registration of histological images of the mouse brain—QuPath^80^ for image segmentation, QuickNII^83^ (RRID: SCR_016854) and VisuAlign (RRID: SCR_017978) for image registration, Qmask for hemispheric differentiation and Nutil^84^ (RRID: SCR_017183) for integration of segmented and registered images. Ten coronal sections per animal which represent the majority of regions with pathology were selected for quantification. To identify pS129 α-syn, individual sections were segmented using the pixel classification feature of QuPath with an optical density threshold of.03 and removal of staining artifacts. The resulting segmentations were exported at full resolution. Down-sampled brain sections were imported into QuickNII and aligned in 3-dimensional space to the 2017 Allen mouse brain atlas CCFv3^82^. Sections were then imported into VisuAlign where anchor points were generated in the atlas and moved to the corresponding location on the section of interest via non-linear warp transformation. In addition, from the spatial coordinate information derived from QuickNII, a custom masking tool Qmask was used to generate masks over each brain hemisphere so ipsilateral and contralateral brain regions could be analyzed independently. The Nutil quantifier feature was then used to integrate the segmentation from QuPath, the registration from VisuAlign, and the hemispheric masks from Qmask to generate percentage area occupied measures for each brain region. The resulting outputs include pixel quantification with spatial location in the brain mapped to 1132 brain regions. All final measures of the percentage of area occupied are reported for each brain region, and statistical analysis is reported for parent regions (e.g. not individual cortical layers).

### Statistical analysis

All statistical analyses other than brain-wide pathology analysis were performed using GraphPad Prism (version 10.2.2). The number of animals analyzed in each experiment, the statistical analysis performed, as well as the p-values for all results < 0.05 are reported in the figure legends. Most statistical analyses were conducted using parametric statistical methods. When comparing vehicle-treated mice (Veh.) to *H. pylori*-infected mice (*H*.*p*.) from the 9-month infection, 3 MPI mPFF, or 6 MPI mPFF cohorts, an F-test was used to compare variances. Those with equal variance were compared via unpaired T-test. Those with unequal variance were compared with Welch’s T-test. When comparing estimated TH^+^ cell counts between hemispheres of 3 MPI mPFF and 6 MPI mPFF cohorts, a two-way ANOVA with uncorrected Fisher’s LSD test was used. When comparing Therapy cohort mice (Veh., *H*.*p*., and *H*.*p*./Ther), One-way ANOVA with Tukey’s multiple comparisons test was used. The data were not log transformed unless otherwise specified in the figure legends. Statistical significance was defined as p < 0.05.

For the analysis assessing differences in brain-wide pathology, data were stratified based on brain hemisphere (Ipsilateral/contralateral), brain region, MPI, and treatment. Regions with zero variance across treatments were excluded from statistical analysis. Significance was determined using second-generation P values based on a null interval of ± 5% difference with 95% confidence intervals^85^. Only second-generation P values equal to 0 were considered significant.

## Supporting information

Supplement for H. pylori manuscript

## ACKNOWLEDGEMENTS

*H. pylori* PMSS1 strain was a gift from Dr. Manuel Amieva (Stanford). We thank the Van Andel Institute Bioinformatics and Biostatistics Core, especially Zachary Madaj and Hannah Damico, for their assistance with statistical analysis, the Van Andel Institute Pathology and Biorepository Core (RRID:SCR_022912) for their assistance with tissue sectioning, the Institute Optical Imaging Core (RRID:SCR_021968) for their maintenance of microscopy equipment, and the Van Andel Institute Vivarium (RRID:SCR_023211) for caring for animals. This research was funded by the Farmer Family Foundation to M.X.H, L. Brundin, and J.A.P, Van Andel Institute, and an Inspire Fellowship to A.S. Several images were created with BioRender.com.

## AUTHOR CONTRIBUTIONS

A.S. designed the experiments, performed experiments, analyzed results and wrote the manuscript. A. Prigent designed experiments and performed experiments. N.S performed experiments. L.M. and L. Breton performed mouse injection surgeries. J.A.P. and L. Brundin provided experimental design guidance and manuscript input, and M.X.H. conceived and designed the experiments and wrote the manuscript. All authors have reviewed and approved this manuscript.

## DATA AND MATERIALS AVAILABILITY

Data and materials are available upon request.

