## Supplement for H. pylori manuscript for "*Helicobacter pylori* infection and α-synuclein pathology drive parallel neurodegenerative pathways in the substantia nigra"

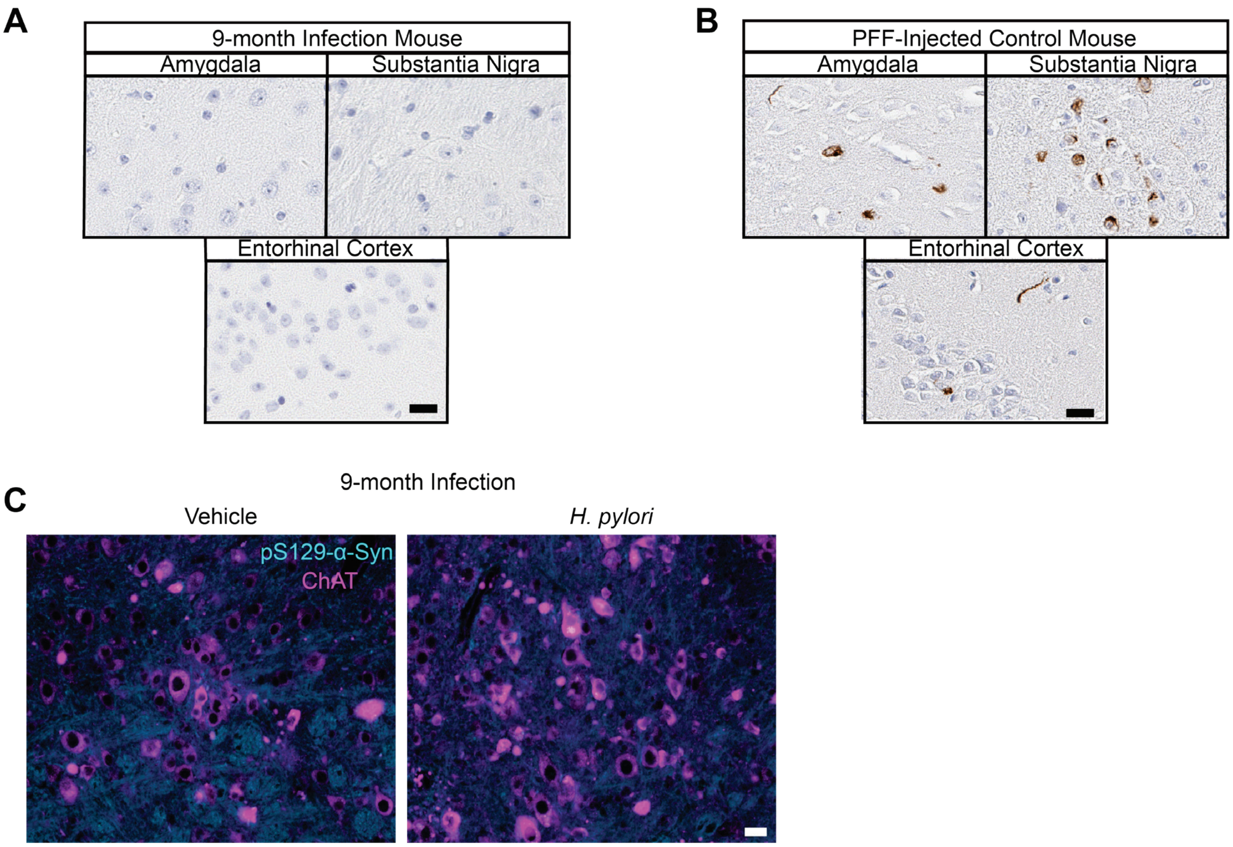


**Fig. S1. Aged *H. pylori*-infected mice do not show signs of α-synucleinopathy in the brain. A.** representative images of *H. pylori*-infected 9-month infection amygdala, substantia nigra, and entorhinal cortex stained for pS129 α-synuclein. **B.** Representative images of α-synuclein PFF-injected positive control amygdala, substantia nigra, and entorhinal cortex stained for pS129 α-synuclein. **C.** Representative image of vehicle-treated and *H. pylori*-infected hindbrains showing the dorsal motor nucleus of the vagus nerve (DMX) stained for choline acetyltransferase (ChAT) and pS129 α-synuclein. The pS129 α-synuclein channel intensity has been increased to demonstrate that the channel was imaged, but only background staining is visible. Scale bars = 20 µm.

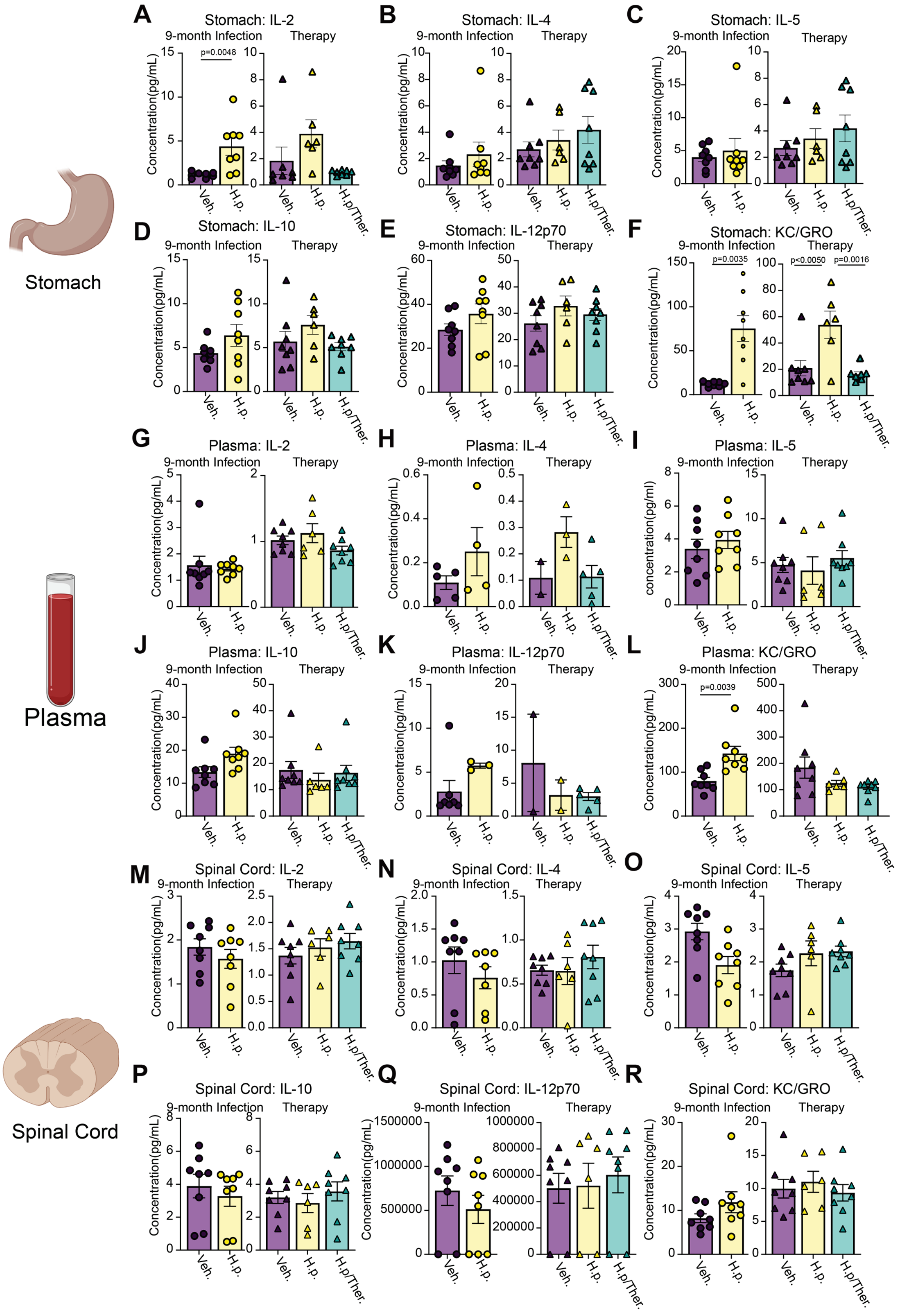

A
Stomach: IL-2
9-month Infection
Therapy
p=0.0048
B
Stomach: IL-4
C
Stomach: IL-5
D
Stomach: IL-10
E
Stomach: IL-12p70
F
Stomach: KC/GRO
p=0.0035
p<0.0050
p=0.0016
Stomach
G
Plasma: IL-2
H
Plasma: IL-4
I
Plasma: IL-5
J
Plasma: IL-10
K
Plasma: IL-12p70
L
Plasma: KC/GRO
p=0.0039
Plasma
M
Spinal Cord: IL-2
N
Spinal Cord: IL-4
O
Spinal Cord: IL-5
P
Spinal Cord: IL-10
Q
Spinal Cord: IL-12p70
R
Spinal Cord: KC/GRO
Spinal Cord
Concentration(pg/mL)
Veh.
H.p.
H.p/Ther.

**Fig. S2. Further evaluation of cytokines and chemokines across tissues.** Multiplexed cytokine assay showing calculated concentration of IL-2, IL-4, IL-5, IL-10, IL-12p70, and KC/GRO levels measured in the **A-F.** stomach tissue, **G-L.** blood plasma, and **M-R.** spinal cord tissue of 9-month infection and Therapy cohort mice. For the 9-month infection cohort, n = 8 (Veh.) or n = 8 (H.p.), Welch's T-tests were performed for **A, C, D, F, G, K,** and **R**. Unpaired T-tests were performed for the remaining panels. For the Therapy cohort, n = 8 (Veh.), 6 (*H.p.)*, or 8 (*H.p.*/Ther.), one-way ANOVA with Tukey's multiple comparisons test was performed. Individual mice with cytokine levels below assay detection limits were excluded from analysis. Statistical significance ($p < 0.05$) is indicated above relevant comparisons; all other differences are non-significant.

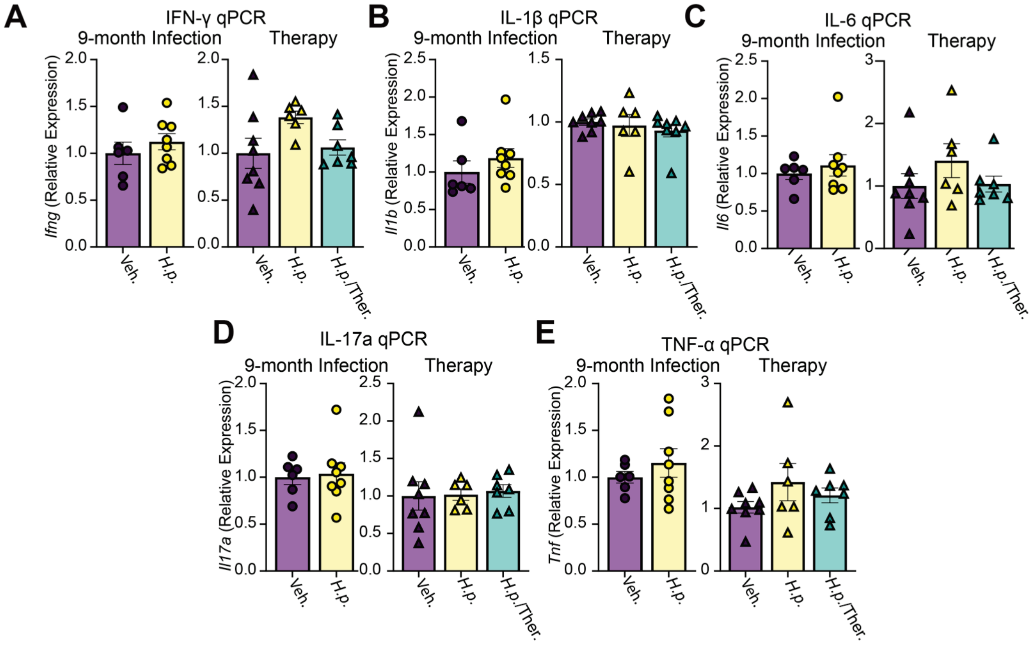


**Fig. S3. mRNA expression of proinflammatory cytokines is not significantly elevated in mouse spinal cords.** Relative expression of **A.** *Ifng*, ***B.*** *Il1b,* ***C.*** *Il6*, **D.** *Il17a*, and **E.** *Tnfa* mRNA in 9-month Infection and Therapy cohort spinal cords. n=6 (Veh.) or 8 (*H.p.)*. Welch's T-test was performed for **E.** Unpaired T-test was performed for **A, B, C,** and **D.** For the Therapy mice, n= 8(Veh.), 6 (*H.p.),* or 7 (*H.p.*/Ther.*).* One-way ANOVA with Tukey's multiple comparisons test was performed. Statistical significance ($p < 0.05$) is indicated above relevant comparisons; all other differences are non-significant.

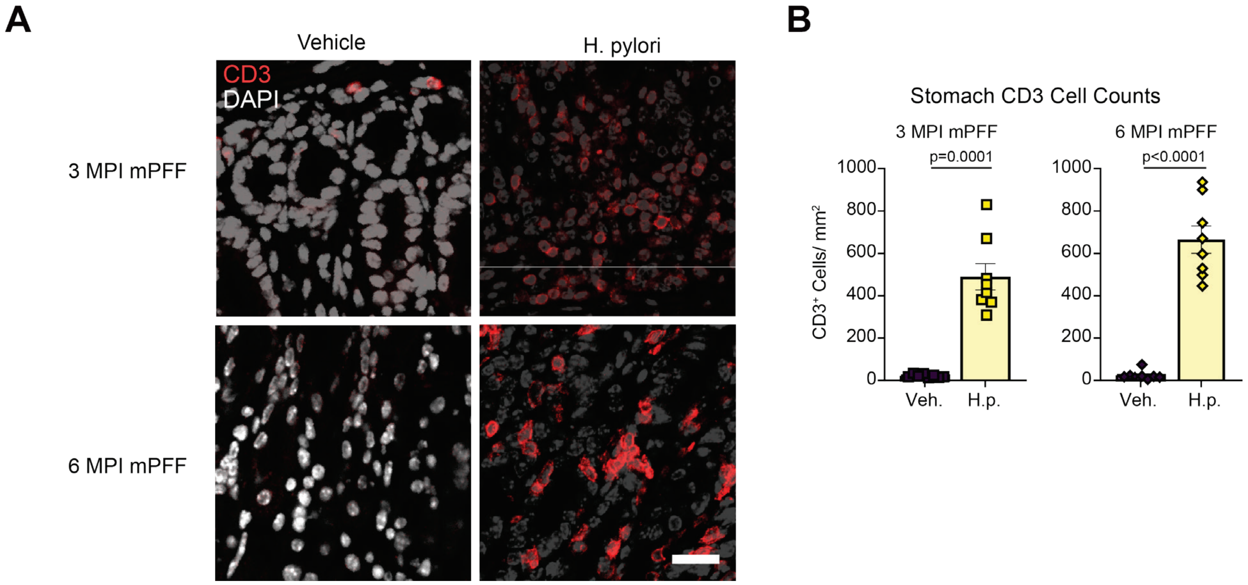


**Fig. S4. Dual-hit *H. pylori* model shows similar prolonged infiltration of the stomach by T-cells. A.** Representative image of 3 MPI PFF and 6 MPI PFF cohort mouse stomachs stained for CD3. Scale bar = 20 µm. **B.** Quantification of $CD3^{+}$ cells. n= 8 (Veh.) or n=8 (*H.p.*). Welch's T-test was performed. Statistical significance ($p < 0.05$) is indicated above relevant comparisons; all other differences are non-significant.

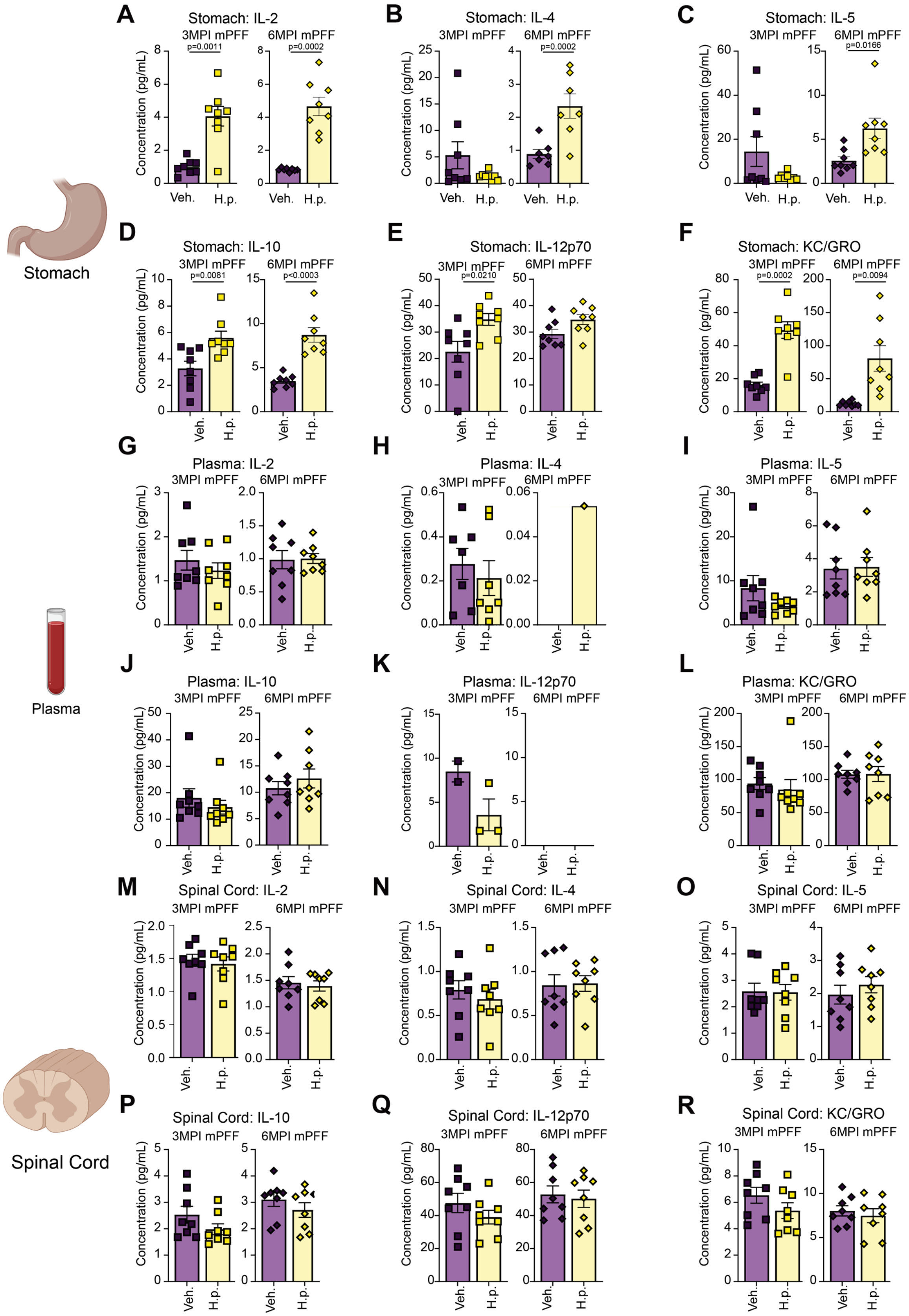

Stomach
Plasma
Spinal Cord
A
Stomach: IL-2
3MPI mPFF
6MPI mPFF
p=0.0011
p=0.0002
B
Stomach: IL-4
3MPI mPFF
6MPI mPFF
p=0.0002
C
Stomach: IL-5
3MPI mPFF
6MPI mPFF
p=0.0166
D
Stomach: IL-10
3MPI mPFF
6MPI mPFF
p=0.0081
p<0.0003
E
Stomach: IL-12p70
3MPI mPFF
6MPI mPFF
p=0.0210
F
Stomach: KC/GRO
3MPI mPFF
6MPI mPFF
p=0.0002
p=0.0094
G
Plasma: IL-2
3MPI mPFF
6MPI mPFF
H
Plasma: IL-4
3MPI mPFF
6MPI mPFF
I
Plasma: IL-5
3MPI mPFF
6MPI mPFF
J
Plasma: IL-10
3MPI mPFF
6MPI mPFF
K
Plasma: IL-12p70
3MPI mPFF
6MPI mPFF
L
Plasma: KC/GRO
3MPI mPFF
6MPI mPFF
M
Spinal Cord: IL-2
3MPI mPFF
6MPI mPFF
N
Spinal Cord: IL-4
3MPI mPFF
6MPI mPFF
O
Spinal Cord: IL-5
3MPI mPFF
6MPI mPFF
P
Spinal Cord: IL-10
3MPI mPFF
6MPI mPFF
Q
Spinal Cord: IL-12p70
3MPI mPFF
6MPI mPFF
R
Spinal Cord: KC/GRO
3MPI mPFF
6MPI mPFF
Concentration (pg/mL)
Veh.
H.p.

**Fig. S5. Further evaluation of cytokines and chemokines in α-synuclein PFF-treated mice.** Multiplexed cytokine assay showing calculated concentration of IL-2, IL-4, IL-5, IL-10, IL-12p70, and KC/GRO levels measured in **A-F.** stomach tissue, **G-L.** blood plasma, and **M-R.** spinal cord tissue of 3 MPI PFF and 6 MPI PFF cohort mice. n = 8 (Veh.) or n = 8 (H.p.), Welch's T-tests were performed for **A, B, C, D,** and **I**. Unpaired T-tests were performed for the remaining panels. Statistical significance ($p < 0.05$) is indicated above relevant comparisons; all other differences are non-significant.

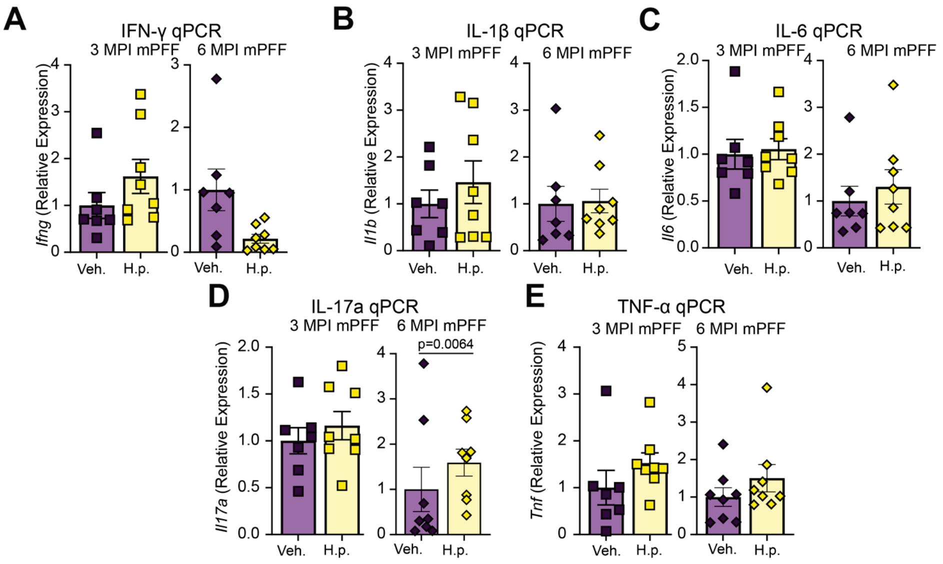


**Fig. S6. Addition of mPFF injections to *H. pylori* infection in mice does not alter expression of proinflammatory cytokines in spinal cords.** Relative expression of **A.** *Ifng*, ***B.*** *Il1b*, ***C.*** *Il6*, **D.** *Il17a*, and **E.** *Tnfa* mRNA in3 MPI mPFF and 6 MPI mPFF spinal cords. n=7 (Veh.) or 8 (*H.p.)*. Welch's T-test was performed for **A** and **D.** Unpaired T-test was performed for **B, C,** and **E.** Statistical significance ($p < 0.05$) is indicated above relevant comparisons; all other differences are non-significant.
